## Supplementary Figures for "Loss of PTPMT1 limits mitochondrial utilization of carbohydrates and leads to muscle atrophy and heart failure in tissue-specific knockout mice"

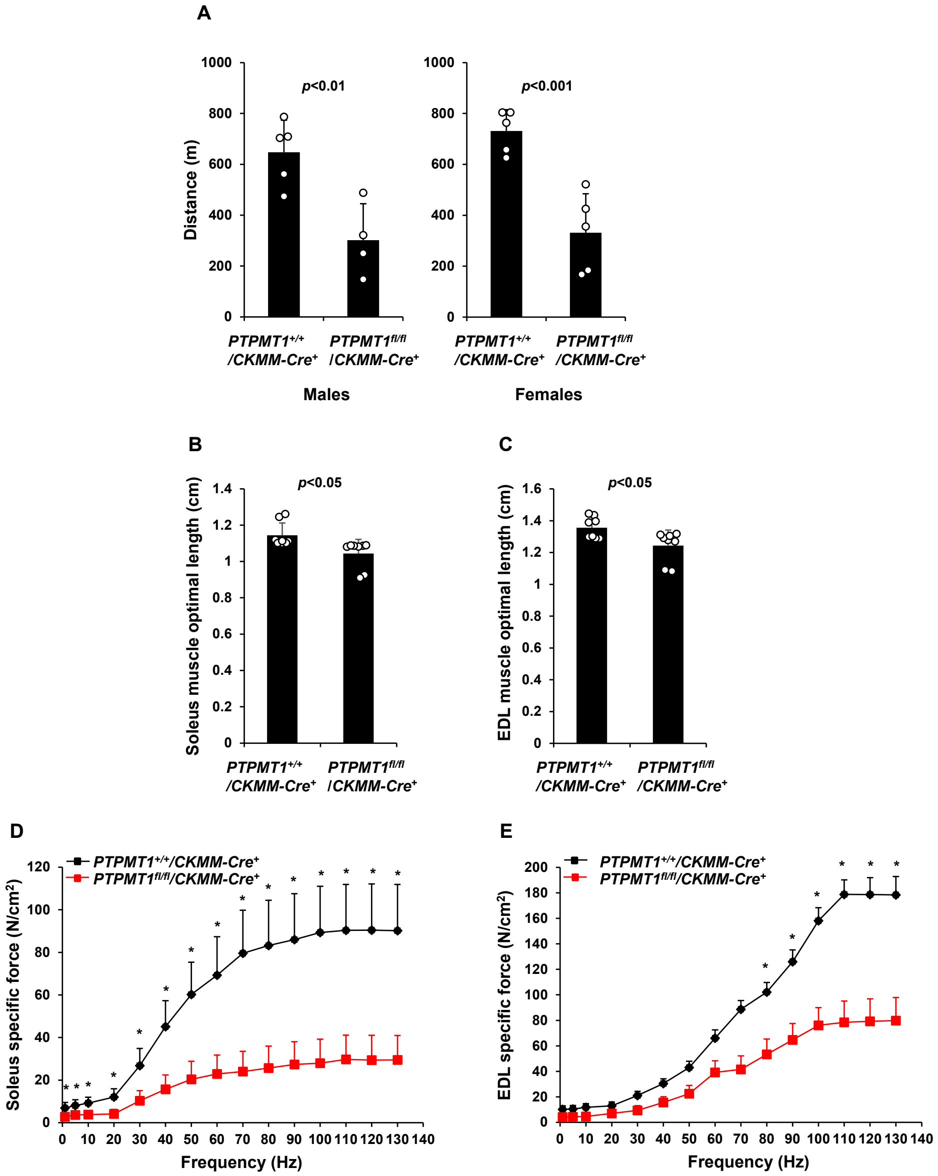


**Fig. S1. Specific force production is decreased in Soleus and EDL isolated from *PTPMT1*$^{fl/fl}$*/CKMM-Cre*$^{+}$ mice.** (**A**) Seven to eight-month-old *PTPMT1*$^{+/+}$*/CKMM-Cre*$^{+}$ (n=5 males and 5 females) and *PTPMT1*$^{fl/fl}$*/CKMM-Cre*$^{+}$ (n=4 males and 5 females) mice were assessed by treadmill exercise tests as described in Materials and Methods. The distance of the run was recorded. (**B** and **C**) Soleus (**B**) and EDL (**C**) dissected from 7 to 8-month-old *PTPMT1*$^{fl/fl}$*/CKMM-Cre*$^{+}$ and *PTPMT1*$^{+/+}$*/CKMM-Cre*$^{+}$ mice (n=4, 8 muscles/genotype) were subjected to length-force relationship tests to determine optimal lengths at which maximal forces were achieved. (**D** and **E**) Soleus (**D**) and EDL (**E**) dissected from 7 to 8-month-old *PTPMT1*$^{fl/fl}$*/CKMM-Cre*$^{+}$ and *PTPMT1*$^{+/+}$*/CKMM-Cre*$^{+}$ mice (n=3, 6 muscles/genotype) were examined by *ex vivo* isometric force measurements. Absolute contractile forces (specific forces) produced at the indicated frequencies of stimulation were documented. * $p < 0.05$.

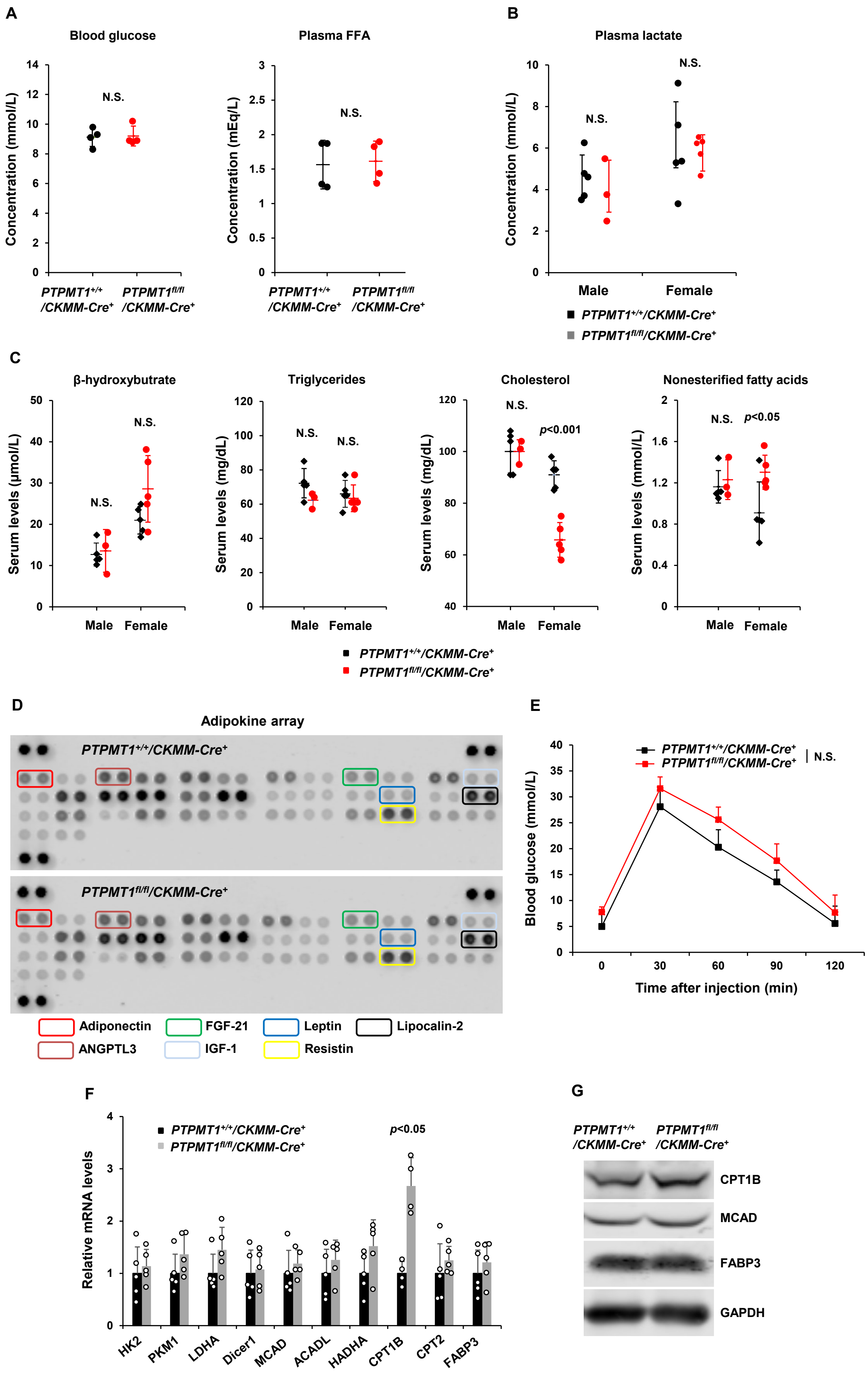
A
Blood glucose
Concentration (mmol/L)
N.S.
PTPMT1+/+ /CKMM-Cre+
PTPMT1fl/fl /CKMM-Cre+
Plasma FFA
Concentration (mEq/L)
N.S.
B
Plasma lactate
Concentration (mmol/L)
N.S.
Male
Female
PTPMT1+/+/CKMM-Cre+
PTPMT1fl/fl/CKMM-Cre+
C
β-hydroxybutrate
Triglycerides
Cholesterol
Nonesterified fatty acids
Serum levels (µmol/L)
Serum levels (mg/dL)
Serum levels (mmol/L)
N.S.
p<0.001
p<0.05
Male
Female
PTPMT1+/+/CKMM-Cre+
PTPMT1fl/fl/CKMM-Cre+
D
Adipokine array
PTPMT1+/+/CKMM-Cre+
PTPMT1fl/fl/CKMM-Cre+
Adiponectin
FGF-21
Leptin
Lipocalin-2
ANGPTL3
IGF-1
Resistin
E
Blood glucose (mmol/L)
PTPMT1+/+/CKMM-Cre+
PTPMT1fl/fl/CKMM-Cre+
N.S.
Time after injection (min)
F
Relative mRNA levels
PTPMT1+/+/CKMM-Cre+
PTPMT1fl/fl/CKMM-Cre+
p<0.05
HK2
PKM1
LDHA
Dicer1
MCAD
ACADL
HADHA
CPT1B
CPT2
FABP3
G
PTPMT1+/+ /CKMM-Cre+
PTPMT1fl/fl /CKMM-Cre+
CPT1B
MCAD
FABP3
GAPDH

**Fig. S2. Plasma levels of glucose and lipids are comparable in *PTPMT1$^{fl/fl}$/CKMM-Cre$^{+}$* mice and *PTPMT1$^{+/+}$/CKMM-Cre$^{+}$* littermates.** (**A**) Blood glucose and free fatty acid (FFA) levels were measured in 6 to 7-month-old male *PTPMT1$^{fl/fl}$/CKMM-Cre$^{+}$* and *PTPMT1$^{+/+}$/CKMM-Cre$^{+}$* mice (n=4/genotype). (**B**) Plasma lactate levels were measured in 6 to 7-month-old *PTPMT1$^{fl/fl}$/CKMM-Cre$^{+}$* and *PTPMT1$^{+/+}$/CKMM-Cre$^{+}$* mice (n=3-5/genotype). (**C**) Serum levels of cholesterol, triglycerides, beta hydroxybutrate, and nonesterified fatty acids in 6 to 7-month-old *PTPMT1$^{fl/fl}$/CKMM-Cre$^{+}$* and *PTPMT1$^{+/+}$/CKMM-Cre$^{+}$* mice (n=5/genotype) were determined. (**D**) Five to seven-month-old *PTPMT1$^{fl/fl}$/CKMM-Cre$^{+}$* and *PTPMT1$^{+/+}$/CKMM-Cre$^{+}$* mice (n=4/genotype) were fasted overnight. Blood samples were collected and 100 µl serum from each mouse was used for the adipokine array assay to determine the relative levels of 38 mouse adipokines with a kit (R&D Systems) following the manufacturer's instructions. Representative images are shown. (**E**) Six to seven-month-old male *PTPMT1$^{fl/fl}$/CKMM-Cre$^{+}$* and *PTPMT1$^{+/+}$/CKMM-Cre$^{+}$* mice (n=8-10/genotype) were deprived of food overnight and then injected intraperitoneally with glucose (2 g/kg). Tail vein blood was sampled for glucose measurements at the indicated time points. (**F** and **G**) Skeletal muscles (Soleus) dissected from 3-month-old *PTPMT1$^{fl/fl}$/CKMM-Cre$^{+}$* and *PTPMT1$^{+/+}$/CKMM-Cre$^{+}$* mice were processed for qRT-PCR (**F**, n=5/genotype) and immunoblotting (**G**, n=3/genotype) assays to determine expression levels of the indicated genes. Representative images are shown.

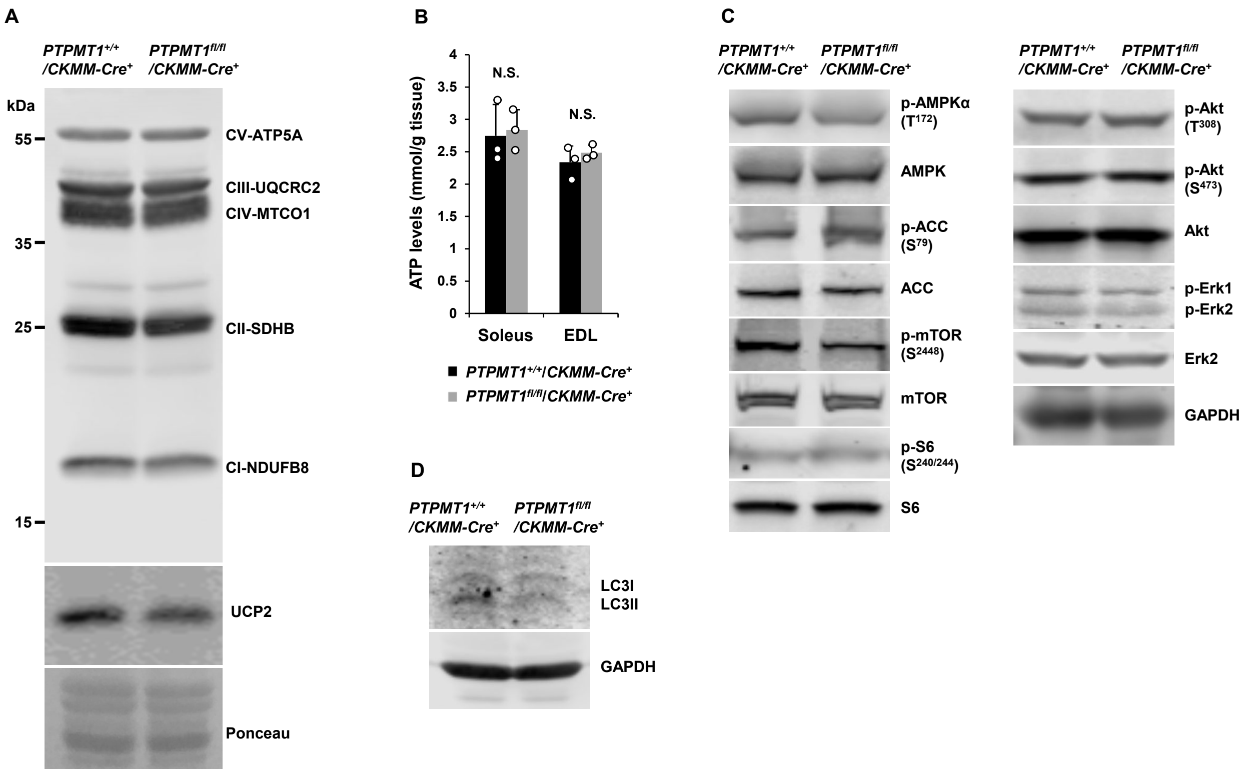


**Fig. S3. Energy homeostasis is initially maintained in young *$PTPMT1^{fl/fl}$/CKMM-$Cre^{+}$* skeletal muscles.** (**A**) Mitochondria were isolated from the skeletal muscles dissected from *$PTPMT1^{fl/fl}$/CKMM-$Cre^{+}$* and *$PTPMT1^{+/+}$/CKMM-$Cre^{+}$* mice (n=3/genotype) at 3 months of age. Expression levels of mitochondrial electron transport chain complexes were determined by immunoblotting. Representative images are shown. (**B**) Total ATP levels in Soleus and EDL dissected from 3-month-old *$PTPMT1^{fl/fl}$/CKMM-$Cre^{+}$* and *$PTPMT1^{+/+}$/CKMM-$Cre^{+}$* mice (n=3/genotype) were determined. (**C** and **D**) Whole cell lysates prepared from Soleus isolated from 3-month-old *$PTPMT1^{fl/fl}$/CKMM-$Cre^{+}$* and control mice were examined by immunoblotting with the indicated antibodies. Representative results (3 mice/genotype) are shown.

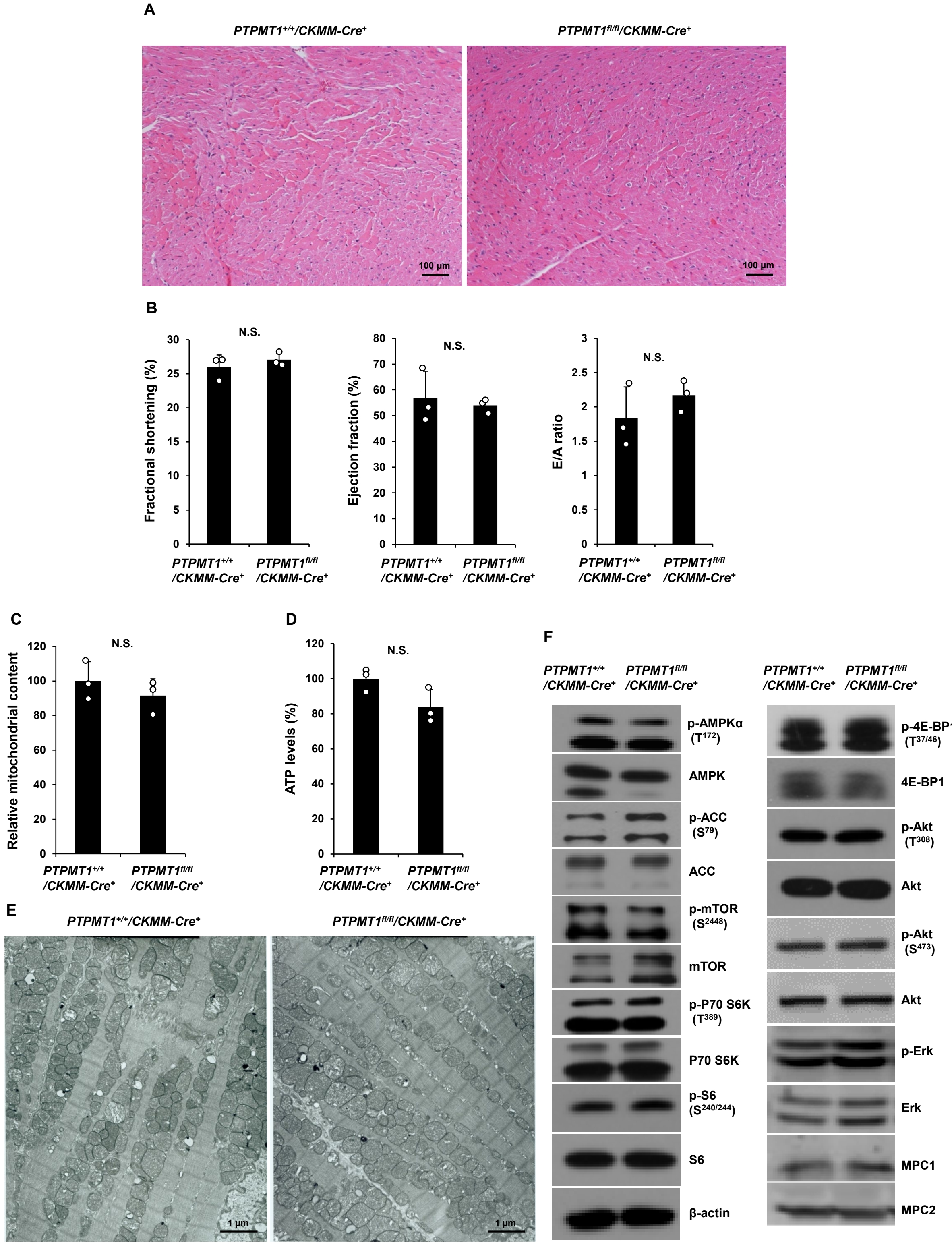
A
PTPMT1+/+/CKMM-Cre+
PTPMT1fl/fl/CKMM-Cre+
100 µm
100 µm
B
N.S.
Fractional shortening (%)
Ejection fraction (%)
E/A ratio
PTPMT1+/+ /CKMM-Cre+
PTPMT1fl/fl /CKMM-Cre+
C
N.S.
Relative mitochondrial content
D
ATP levels (%)
E
1 µm
F
p-AMPKα (T172)
AMPK
p-ACC (S79)
ACC
p-mTOR (S2448)
mTOR
p-P70 S6K (T389)
P70 S6K
p-S6 (S240/244)
S6
β-actin
p-4E-BP1 (T37/46)
4E-BP1
p-Akt (T308)
Akt
p-Akt (S473)
Akt
p-Erk
Erk
MPC1
MPC2

**Fig. S4. Heart functions are normal in *PTPMT1$^{fl/fl}$/CKMM-Cre$^{+}$* mice at 7 months.** (**A**) Heart tissue sections prepared from 7-month-old *PTPMT1$^{+/+}$/CKMM-Cre$^{+}$* and *PTPMT1$^{fl/fl}$/CKMM-Cre$^{+}$* mice were processed for H&E staining. Representative images (3 mice/genotype) are shown. (**B**) Cardiac functions of 5-month-old *PTPMT1$^{+/+}$/CKMM-Cre$^{+}$* and *PTPMT1$^{fl/fl}$/CKMM-Cre$^{+}$* mice (n=3/genotype) were examined by echocardiography. LV fractional shortening (FS), ejection fraction (EF), and ratios of peak velocity of early to late filling of mitral inflow (E/A) were determined. (**C**) Total DNA was extracted from the heart tissues isolated from *PTPMT1$^{fl/fl}$/CKMM-Cre$^{+}$* and *PTPMT1$^{+/+}$/CKMM-Cre$^{+}$* mice (n=3/genotype). Mitochondrial content was estimated by comparing mtDNA (Cytochrome B) levels to genomic DNA levels by quantitative PCR. (**D**) Total ATP levels in the heart tissues isolated from 8-month-old *PTPMT1$^{fl/fl}$/CKMM-Cre$^{+}$* and *PTPMT1$^{+/+}$/CKMM-Cre$^{+}$* mice (n=3/genotype) were determined. (**E**) Heart tissues dissected from 7-month-old *PTPMT1$^{fl/fl}$/CKMM-Cre$^{+}$* and *PTPMT1$^{+/+}$/CKMM-Cre$^{+}$* mice were processed for transmission electron microscopy examination. Representative images (3 mice/genotype) are shown. (**F**) Whole cell lysates prepared from the heart tissues dissected from 7-month-old *PTPMT1$^{fl/fl}$/CKMM-Cre$^{+}$* and *PTPMT1$^{+/+}$/CKMM-Cre$^{+}$* mice were examined by immunoblotting with the indicated antibodies. Representative results (3 mice/genotype) are shown.

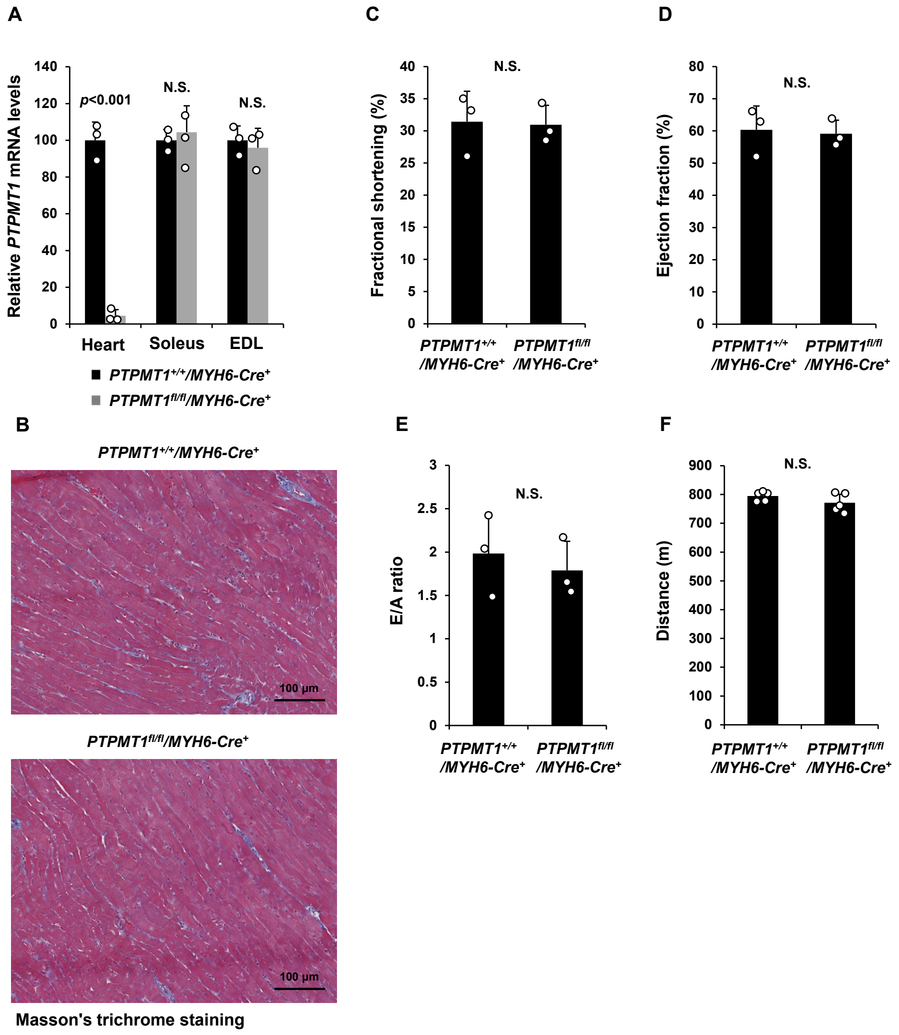


**Fig. S5. Cardiac functions are normal in middle-aged or younger heart-specific *PTPMT1* knockout mice.** (**A**) *PTPMT1* mRNA levels in the heart, Soleus, and EDL dissected from *PTPMT1*$^{+/+}$*/MYH6-Cre*$^{+}$ and *PTPMT1*$^{fl/fl}$*/MYH6-Cre*$^{+}$ mice (n=3/genotype) were determined by qRT-PCR. (**B**) Heart tissue sections prepared from 3-month-old *PTPMT1*$^{fl/fl}$*/MYH6-Cre*$^{+}$ and *PTPMT1*$^{+/+}$*/MYH6-Cre*$^{+}$ mice were processed for Masson's Trichrome staining. Representative images (3 mice/genotype) are shown. (**C**-**E**) Echocardiographic analyses were performed for 3-month-old *PTPMT1*$^{fl/fl}$*/MYH6-Cre*$^{+}$ and *PTPMT1*$^{+/+}$*/MYH6-Cre*$^{+}$ mice (n=3/genotype). LV FS (**C**), EF (**D**), and E/A ratios (**E**) were determined. (**F**) Six-month-old *PTPMT1*$^{fl/fl}$*/MYH6-Cre*$^{+}$ and *PTPMT1*$^{+/+}$*/MYH6-Cre*$^{+}$ male mice (n=5/genotype) were subjected to treadmill exercise tests. Total distances traveled before the mice collapsed were recorded.

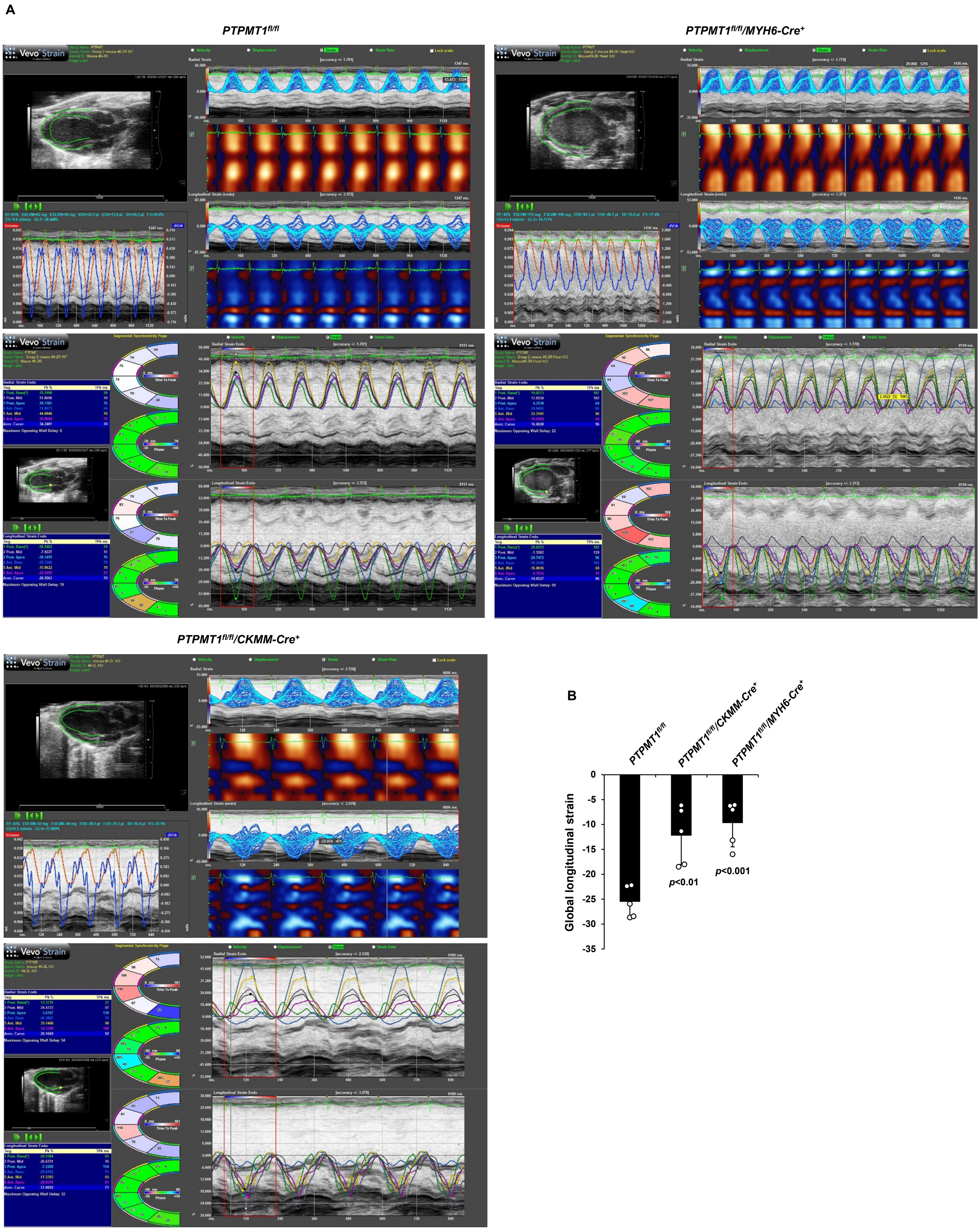
A
PTPMT1fl/fl
PTPMT1fl/fl/MYH6-Cre+
PTPMT1fl/fl/CKMM-Cre+
B
PTPMT1fl/fl
PTPMT1fl/fl/CKMM-Cre+
PTPMT1fl/fl/MYH6-Cre+
Global longitudinal strain
0
-5
-10
-15
-20
-25
-30
-35
p<0.01
p<0.001

**Fig. S6. Cardiac dysfunction in 12-month-old *PTPMT1$^{fl/fl}$/CKMM-Cre$^{+}$* and *PTPMT1$^{fl/fl}$/MYH6-Cre$^{+}$* mice.** (**A** and **B**) Echocardiographic speckle-tracking-based strain measurements were conducted for 12-month-old *PTPMT1$^{fl/fl}$, PTPMT1$^{fl/fl}$/CKMM-Cre$^{+}$*, and *PTPMT1$^{fl/fl}$/MYH6-Cre$^{+}$* mice (n=5/genotype). Representative strain images are shown (**A**). LV global longitudinal strain (GLS) was determined (**B**).

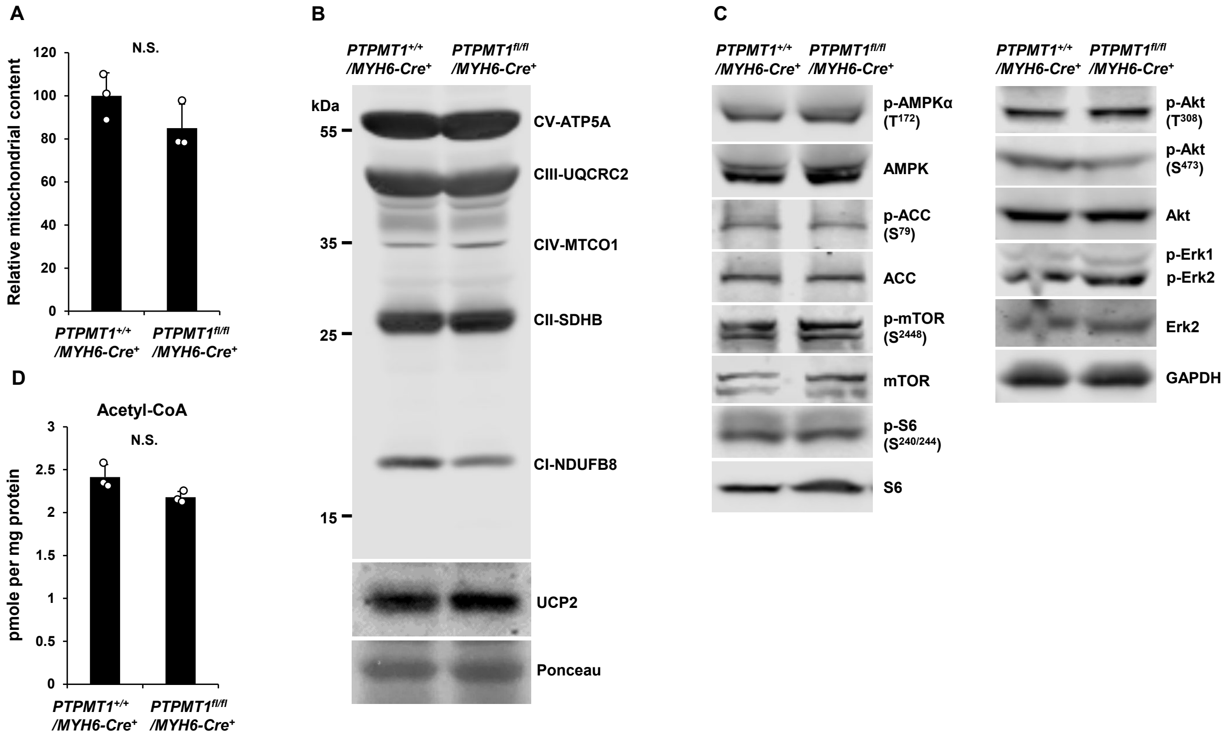


**Fig. S7. There is no bioenergetic stress in young *PTPMT1* knockout cardiomyocytes.** (**A**) Total DNA was extracted from the heart tissues isolated from *PTPMT1$^{fl/fl}$/MYH6-Cre$^{+}$* and *PTPMT1$^{+/+}$/MYH6-Cre$^{+}$* mice (n=3/genotype). Mitochondrial content was estimated by comparing mtDNA (Cytochrome B) levels to genomic DNA levels by quantitative PCR. (**B**) Mitochondria were isolated from the heart tissues freshly dissected from 3-month-old *PTPMT1$^{fl/fl}$/MYH6-Cre$^{+}$* and *PTPMT1$^{+/+}$/MYH6-Cre$^{+}$* mice (n=3/genotype). Expression levels of mitochondrial electron transport chain complexes were determined by immunoblotting. Representative images are shown. (**C**) Whole cell lysates prepared from the heart tissues dissected from 3-month-old *PTPMT1$^{fl/fl}$/MYH6-Cre$^{+}$* and *PTPMT1$^{+/+}$/MYH6-Cre$^{+}$* mice were examined by immunoblotting with the indicated antibodies. Representative results (3 mice/genotype) are shown. (**D**) Mitochondria were isolated from the heart tissues dissected from 2 to 3-month-old *PTPMT1$^{fl/fl}$/MYH6-Cre$^{+}$* and *PTPMT1$^{+/+}$/MYH6-Cre$^{+}$* mice (n=3 mice/genotype). Acetyl-CoA levels in the mitochondrial lysates were determined using an acetyl-CoA assay kit following the manufacturer's instructions.

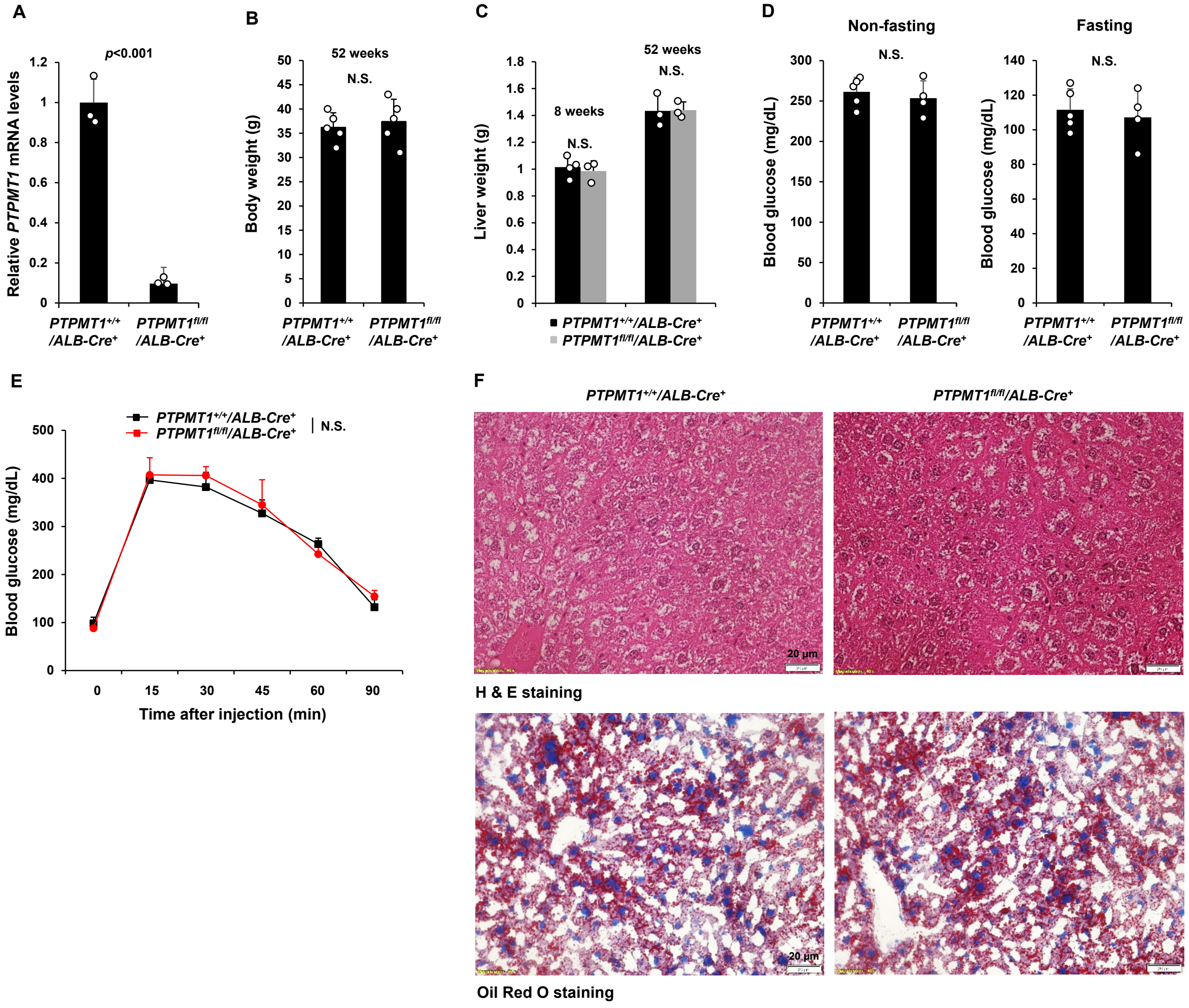


**Fig. S8. The impact of *PTPMT1* deletion from the liver is minimal.** (**A**) *PTPMT1* mRNA levels in the livers dissected from *PTPMT1*$^{+/+}$*/ALB-Cre*$^{+}$ and *PTPMT1*$^{fl/fl}$*/ALB-Cre*$^{+}$ mice (n=3/genotype) were determined by qRT-PCR. (**B** and **C**) Body weights (n=5 mice/genotype) (**B**) and liver weights (n=3-4 mice/genotype) (**C**) of *PTPMT1*$^{+/+}$*/ALB-Cre*$^{+}$ and *PTPMT1*$^{fl/fl}$*/ALB-Cre*$^{+}$ male mice at the indicated ages were determined. (**D**) Blood glucose levels were measured in 8-week-old *PTPMT1*$^{fl/fl}$*/ALB-Cre*$^{+}$ and *PTPMT1*$^{+/+}$*/ALB-Cre*$^{+}$ male mice (n=4-5/genotype) under feeding and overnight-fasting conditions. (**E**) Eight-week-old *PTPMT1*$^{fl/fl}$*/ALB-Cre*$^{+}$ and *PTPMT1*$^{+/+}$*/ALB-Cre*$^{+}$ mice (n=5/genotype) were deprived of food overnight and then injected intraperitoneally with glucose (2 g/kg). Tail vein blood was sampled for glucose measurements at the indicated time points. (**F**) Liver sections prepared from 12-month-old *PTPMT1*$^{fl/fl}$*/ALB-Cre*$^{+}$ and *PTPMT1*$^{+/+}$*/ALB-Cre*$^{+}$ mice were processed for H&E and Oil Red O staining. Representative pictures (3 mice/genotype) are shown.

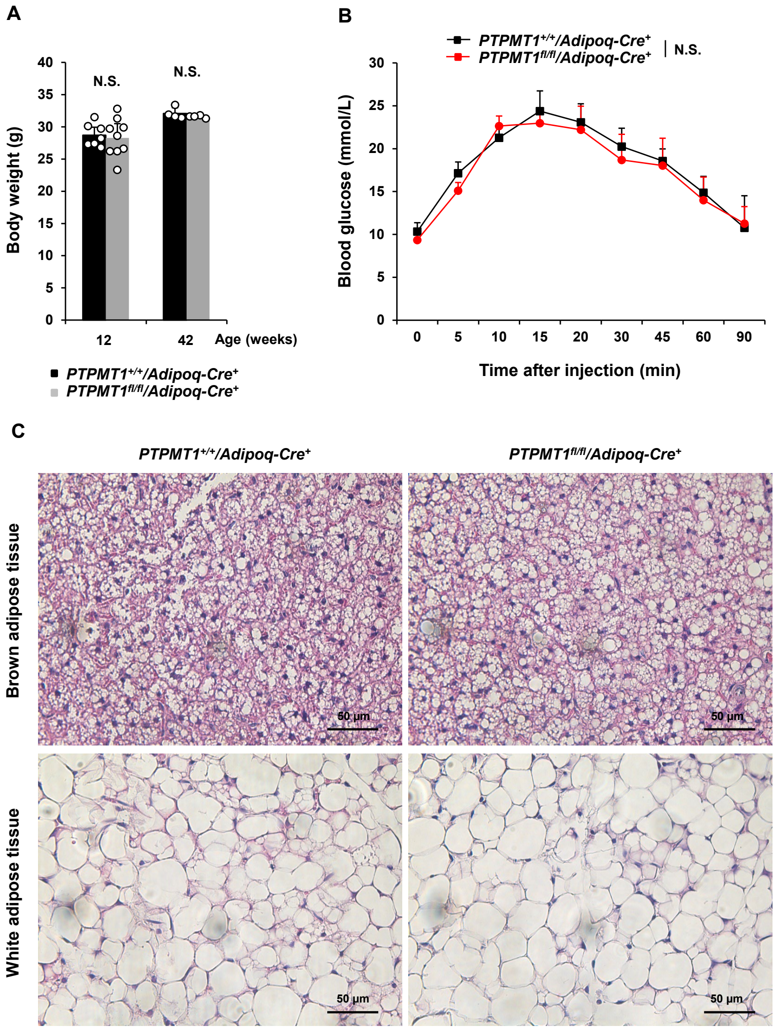


**Fig. S9. No defects are observed in adipocyte-specific *PTPMT1* knockout mice.** (**A**) Body weights of *PTPMT1*$^{fl/fl}$*/Adipoq-Cre*$^{+}$ and *PTPMT1*$^{+/+}$*/Adipoq-Cre*$^{+}$ mice of 12 weeks (n=7-9/genotype) and 42 weeks (n=4/genotype) were determined. (**B**) Eight-week-old *PTPMT1*$^{fl/fl}$*/Adipoq-Cre*$^{+}$ (n=5) and *PTPMT1*$^{+/+}$*/Adipoq-Cre*$^{+}$ (n=4) mice were deprived of food overnight and then injected intraperitoneally with glucose (2 g/kg). Tail vein blood was sampled for glucose measurements at the indicated time points. (**C**) Brown adipose and white adipose tissue sections prepared from 6-week-old *PTPMT1*$^{fl/fl}$*/Adipoq-Cre*$^{+}$ and *PTPMT1*$^{+/+}$*/Adipoq-Cre*$^{+}$ mice were processed for H&E staining. Representative pictures (3 mice/genotype) are shown.

**Full unedited blots images for Figure 2F**

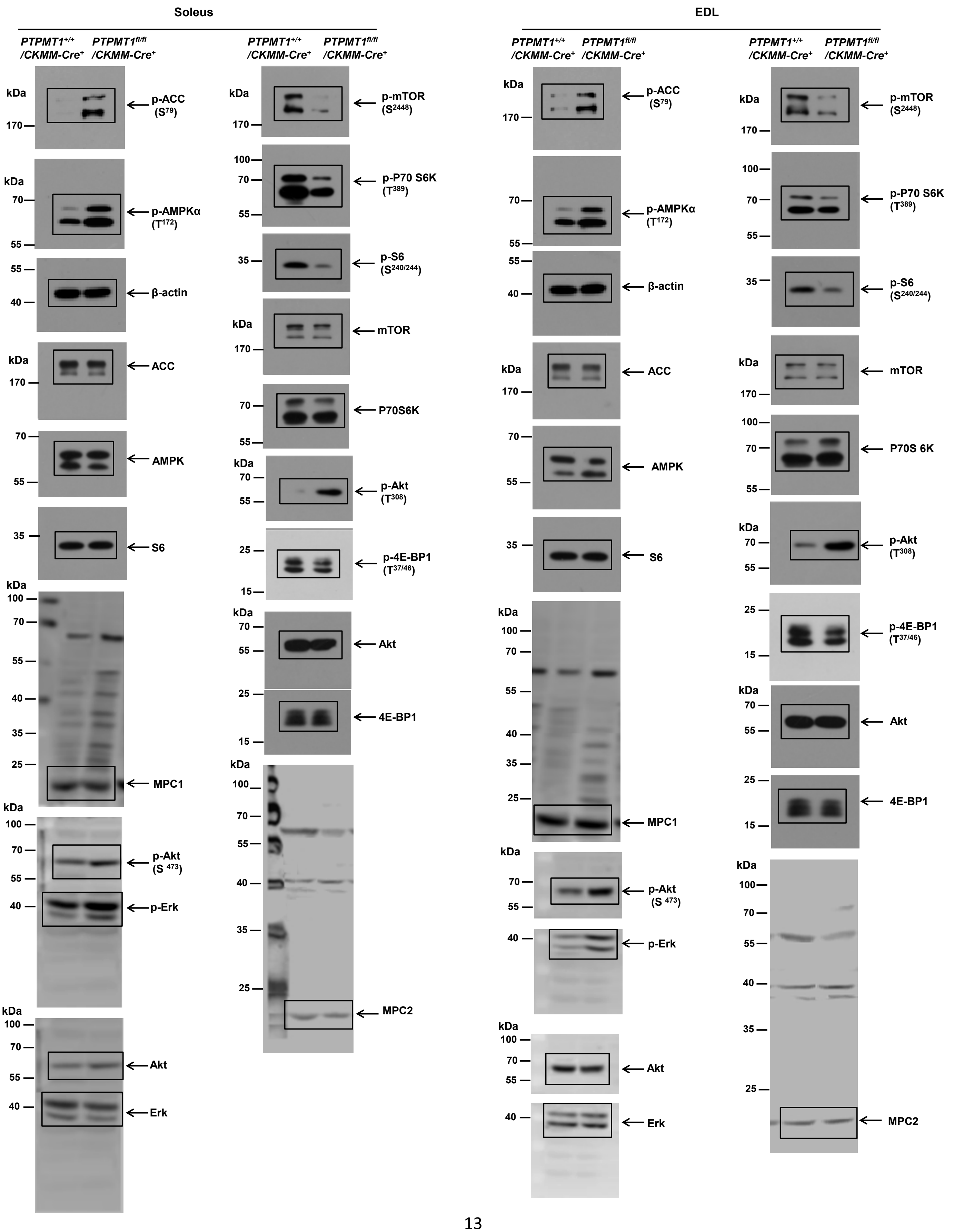

**Full unedited blots images for Figure 6B**

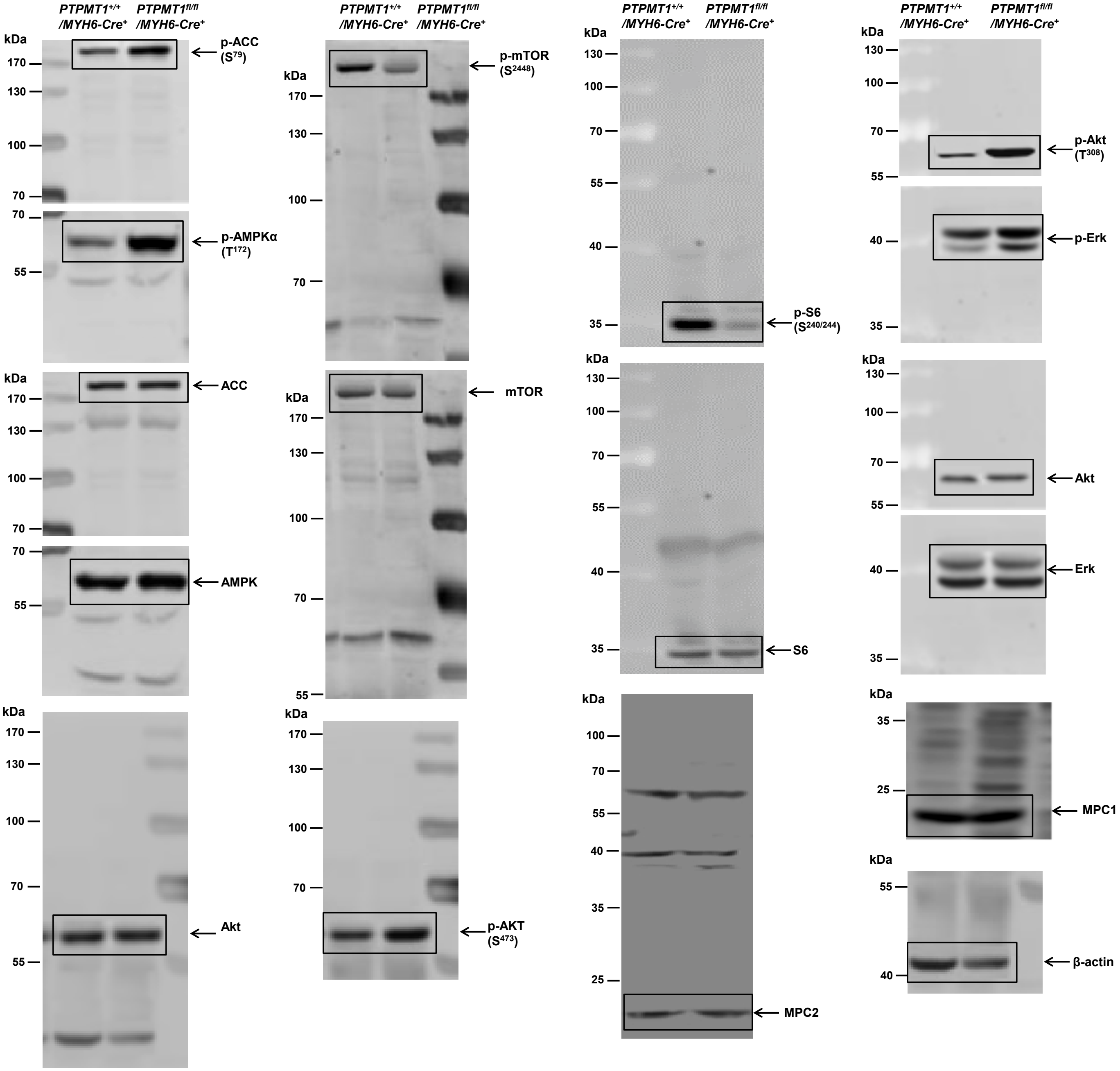

**Full unedited blots images for Supplemental Figure 2G**

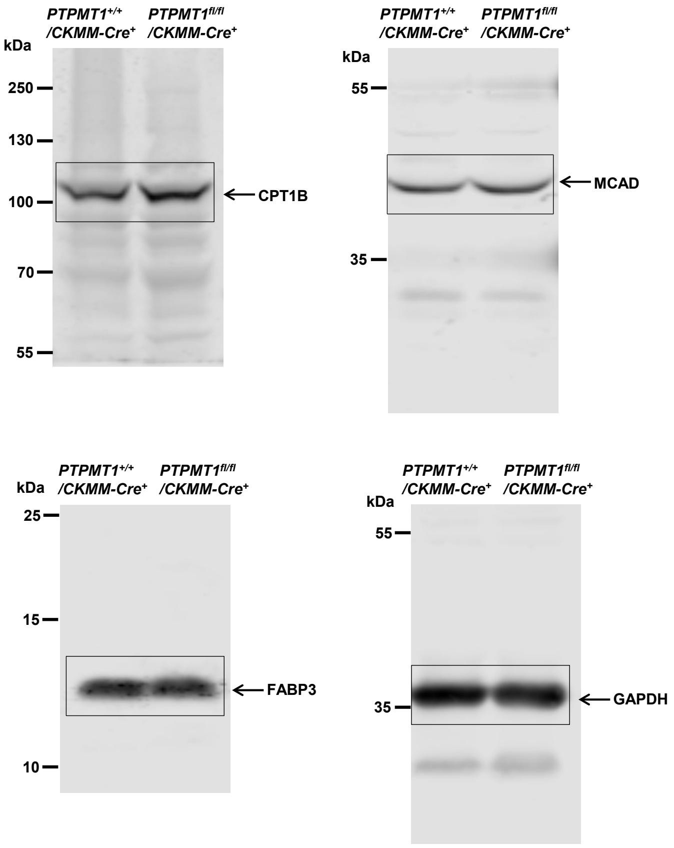

**Full unedited blots images for Supplemental Figure 3C and 3D**

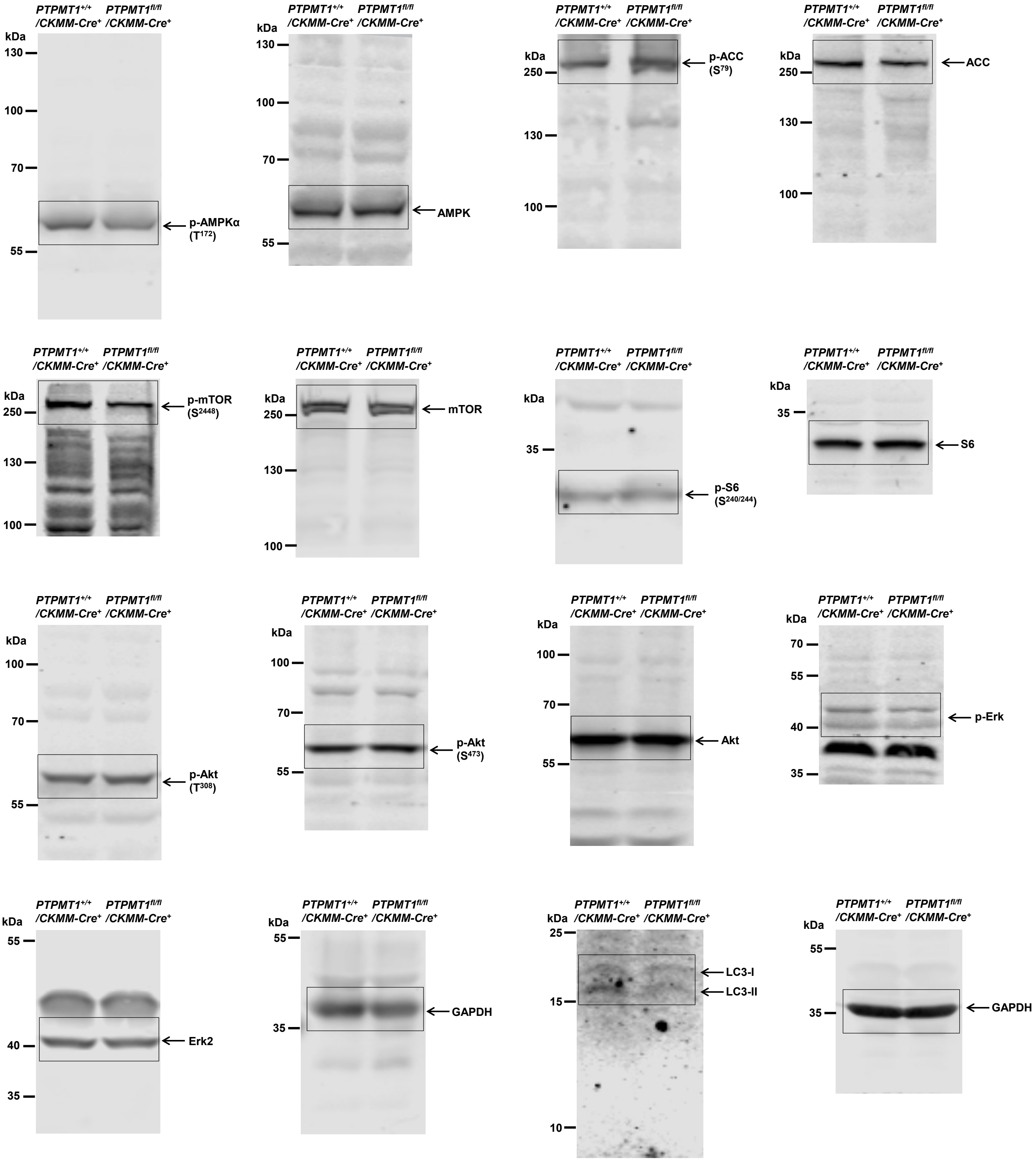

**Full unedited blots images for Supplemental Figure 4F**

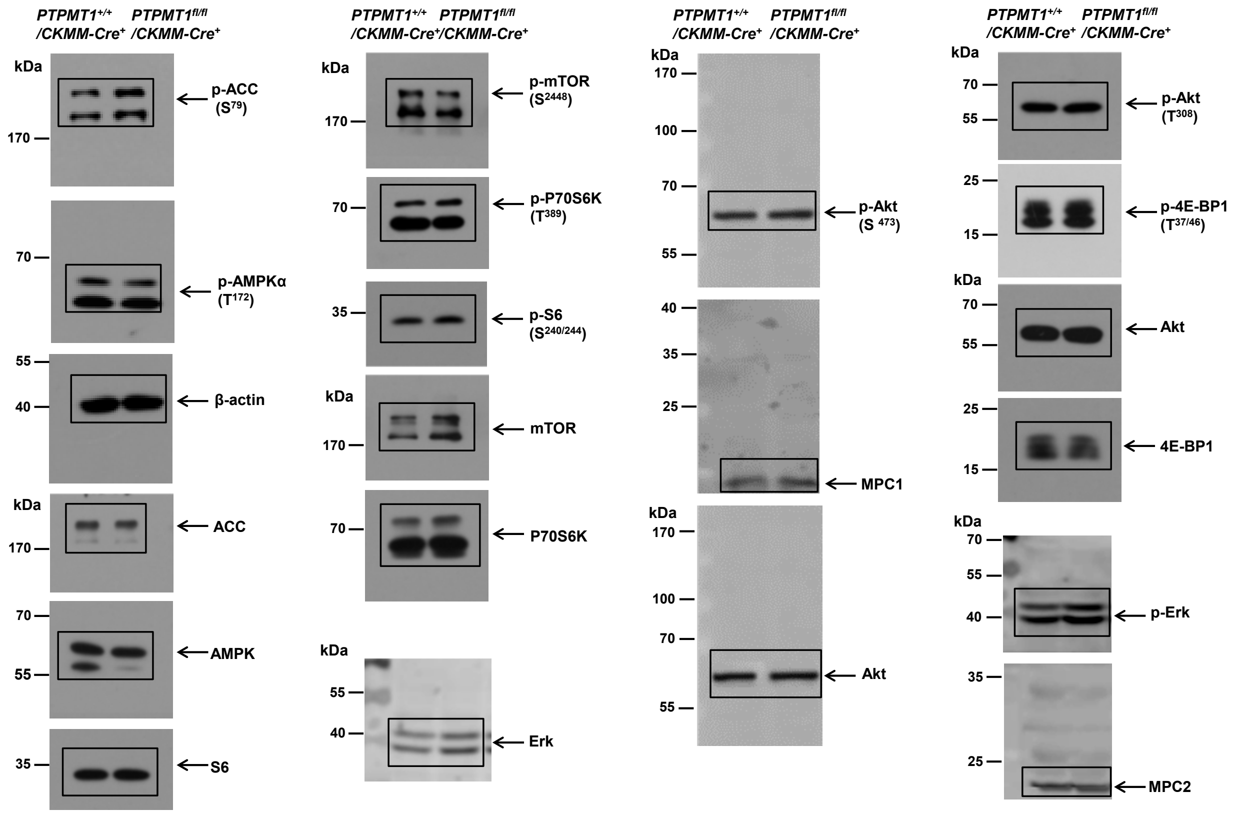

**Full unedited blots images for Supplemental Figure 7C**

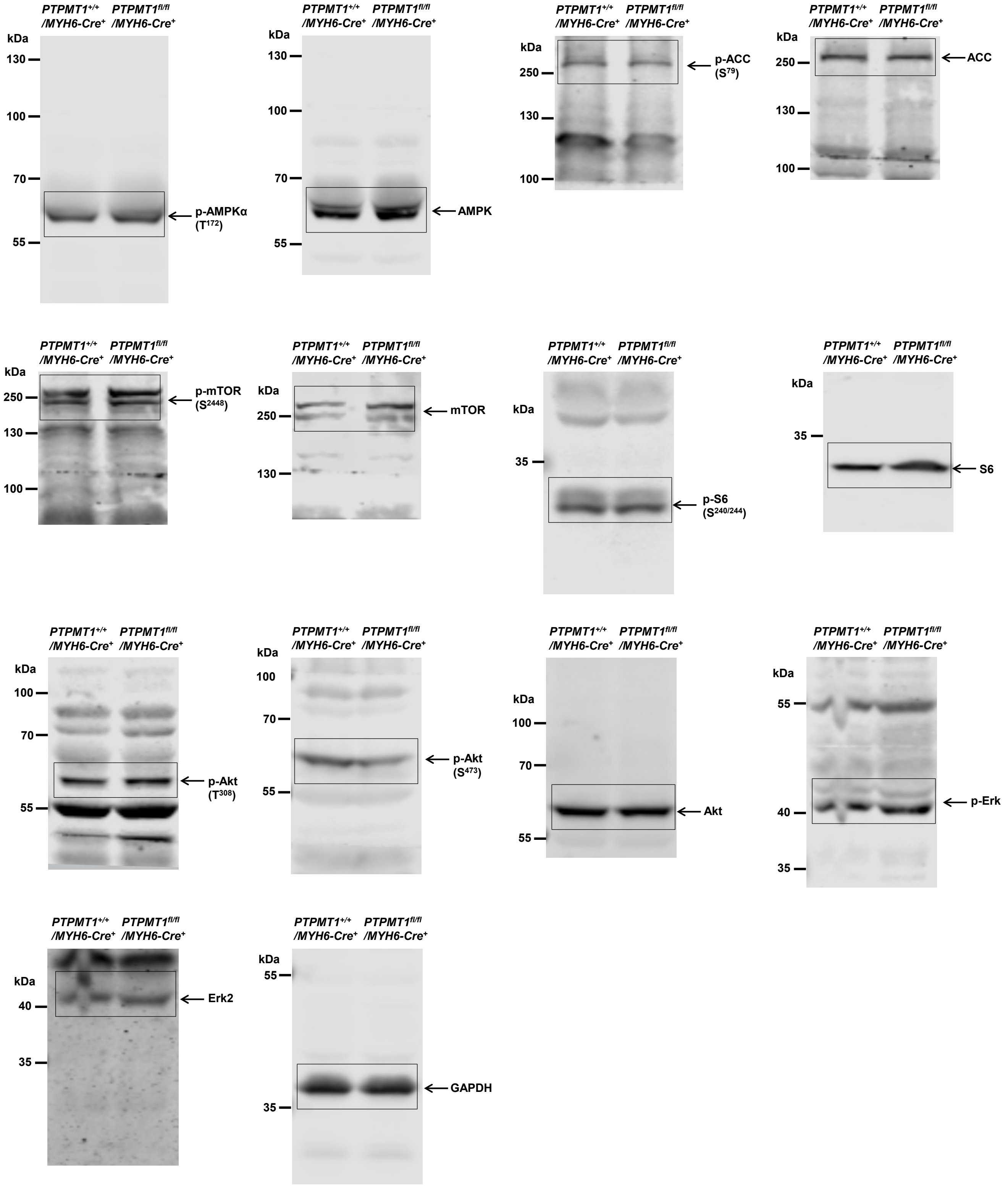

**Fig. S10. Uncropped immunoblotting images.** Uncropped immunoblotting images of Figures 2F and 6B, Supplemental Figures 2G, 3C, 3D, 4F, and 7C are shown. The protein molecular weight markers used were PageRuler™ Prestained Protein Ladder (Thermo Fisher Scientific) and PageRuler™ Prestained Protein Ladder (Fermentas). KDa = kilodalton.
