## Supplementary Video Legends for "Loss of PTPMT1 limits mitochondrial utilization of carbohydrates and leads to muscle atrophy and heart failure in tissue-specific knockout mice"

**Video S1. Cardiac dysfunction in *PTPMT1$^{fl/fl}$/CKMM-Cre$^{+}$* mice.** Twelve-month-old *PTPMT1$^{fl/fl}$/CKMM-Cre$^{+}$* (**A**) and *PTPMT1$^{+/+}$/CKMM-Cre$^{+}$* (**B**) mice (n=5/genotype) were anesthetized with 2% isoflurane and 2-D echocardiography was performed using the Vevo 2100 system and MS400 transducer. Representative 2-D (B-mode) long-axis recordings of left ventricular function are shown.

**Video S2. Heart failure in *PTPMT1$^{fl/fl}$/MYH6-Cre$^{+}$* mice.** Eleven to twelve-month-old *PTPMT1$^{fl/fl}$/MYH6-Cre$^{+}$* (**A**) and *PTPMT1$^{+/+}$/MYH6-Cre$^{+}$* (**B**) mice (n=5/genotype) were anesthetized with 2% isoflurane and 2-D echocardiography was performed using the Vevo 2100 system and MS400 transducer. Representative 2-D (B-mode) long-axis recordings of left ventricular function are shown.
